# *Sporaefaciens musculi* gen. nov., sp. nov., a novel bacterium isolated from the caecum of an obese mouse

**DOI:** 10.1101/2019.12.21.885665

**Authors:** Torben Sølbeck Rasmussen, Theresa Streidl, Thomas C.A. Hitch, Esther Wortmann, Paulina Deptula, Michael Hansen, Dennis Sandris Nielsen, Thomas Clavel, Finn Kvist Vogensen

**Author notes:** Correspondence: Torben S. Rasmussen – & Finn K. Vogensen.

## Abstract

A bacterial strain, designated WCA-9-b2, was isolated from the caecal content of an 18-week-old obese C57BL/6NTac male mouse. According to phenotypic analyses, the isolate is rod-shaped, Gram-positive, strictly anaerobic, spore-forming and non-motile under the conditions tested. Bacterial colonies were irregular and non-pigmented. Analysis of the 16S rRNA gene indicated that the isolate belonged to the family *Lachnospiraceae* with *Clostridium scindens* ATTC 35704 (94.9% sequence identity) and *Dorea formicigenerans* ATCC 27755 (94.8%) being the closest relatives. Whole genome sequencing showed average nucleotide identity (ANI) ranging from 69.80–74.23% and percentage of conserved proteins (POCP) values < 50% against the nine closest relatives. The genome-based G+C content of genomic DNA was 44.4%. The predominant metabolic end products of glucose fermentation were acetate and succinate. Based on these data, we propose that strain WCA-9-b2 represents a novel species within a novel genus, for which the name *Sporaefaciens musculi* gen. nov., sp. nov. is proposed. The type strain is WCA-9-b2^T^ (=DSM 106039^T^ = CCUG pending ID^T^).

**Repositories:** The GenBank accession number for the 16S rRNA gene sequence of strain WCA-9-b2^T^ is MN756014, and the accession number for the genome assembly is PRJNA592877. Raw sequencing Illumina NextSeq (PRJEB35655) and ONT MinION (PRJEB35656) data can be accessed at EMBL-EBI.

## Introduction

The mammalian gut is inhabited by a high diversity of strictly anaerobic bacteria predominated by Gram-positive species from the phylum *Firmicute*s [1], of which many have not yet been cultured. Metagenome sequencing has been an efficient tool to perform *in silico* characterisation of the unculturable gut bacteria [2], although metagenome sequencing often misses spore-forming bacteria due to the difficulty of DNA extraction from the robust structure of the spores [3]. As the mammalian gut microbiota (GM) influences host health [4–6], combining *in silico* characterisation of gut bacteria with culturing novel isolates *in vitro* opens new research avenues by investigating their functional properties. A large diversity of bacteria belonging to the class *Clostridia* contribute in maintaining gut health, amongst other through the production of short chain fatty acids (SCFAs) [7, 8]. Especially, butyrate has been associated with beneficial health effects [7], even though many butyrate producers are dependent on cross-feeding from other bacteria producing intermediate products such as acetate or lactate [9]. Whereas succinate, an intermediate product of propionate, is a double-edge sword that in some conditions sustain the stability of the GM component by cross-feeding, whilst in other conditions elevates the pathogenic potential of infectious bacteria such as *Clostridioides difficile* or *Clostridium rodentium.* In the current study we report the cultivation and detailed characterisation of a spore-forming, succinate and acetate producing bacterial strain WCA-9-b2^T^ isolated from the gut of an obese male mouse, which we propose represents a novel genus within family *Lachnospiraceae*, phylum *Firmicutes*.

## Isolation and ecology

Strain WCA-9-b2^T^ originated from the caecum of a C57BL/6NTac mouse fed a high-fat diet (HF, Research Diets D12492, USA) *ad libitum* for 13 weeks as part of a previously published study [10]. The initial isolation of strain WCA-9-b2 took place at the ZIEL Core Facility Microbiome of the Technical University of Munich (TUM), Germany. Downstream analyses were conducted at the University of Copenhagen (UCPH), Denmark. Anaerobic handling and agar plate incubation during isolation at TUM was performed in an anaerobic chamber (VA500 workstation, Don Whitley Scientific) containing an atmosphere of 90 % (v/v) N_2_ and 10 % H_2_ a at a temperature of 37°C and a humidity of 75 %. All materials were brought into the anaerobic chamber at least 24 hours before use to ensure anaerobic conditions. Liquid and solid media were autoclaved and contained 0.02% 1,4-Dithiothreitol (Sigma, cat. No. DTT-RO) and 0.05% L-cysteine (Sigma, cat. no. 168149) as reducing agents and 1 mg/L resazurin as a redox potential indicator. Broth media were heated prior to flushing with 100% N_2_ by an anaerobic gassing unit (AGU, QCAL Messtechnik GmbH) for at least 3 min. pr. 10 mL. Agar media in Petri dishes contained 1.5% agar (Oxoid™, cat. no. LP0011). Incubations took place at 37°C unless otherwise stated.

Strain WCA-9-b2^T^ was isolated on Wilkens-Chalgren-Anaerobe agar (WCA, Sigma, cat. no. W1886) supplemented with 0.1% bile salt (WCA-BS) (sodium taurocholate hydrate, Sigma, cat. no. 86339) to enhance spore germination. Approx. ∼20 mg thawed caecum content was transferred to a 0.20 µm sterile filtered 1:1 mixture of PBS (NaCl 137mM, KCl 2.7 mM, Na_2_HPO_4_ 10 mM, KH_2_PO_4_ 1.8mM) and 70% ethanol to kill vegetative cells. The suspension was incubated at RT (room temperature) under aerobic conditions for four hours and vortexed every 30 min., followed by 3x centrifugation (11000 x g, 5 min.) and resuspension in aerobic PBS. Ten-fold serial dilutions were prepared with anoxic PBS containing 0.02 % peptone (Sigma, cat. no. 91249). Each dilution (10 µl) was deposited on WCA-BS agar and immediately tilted for vertical migration on the plate, followed by incubation at 37°C for 4 days. Bacterial colonies from a representative area were picked and further re-streaked on WCA agar 3x to obtain pure cultures. WCA-9-b2^T^ was deposited at the Deutsche Sammlung von Mikroorganismen und Zellkulturen GmbH (DSM 106039) and the Culture Collection University of Gothenburg (CCUG pending ID^T^). Liquid nitrogen was used to snap-freeze cryo-aliquots (pivotal for survival of the strain), i.e., cultures diluted 1:1 in 50% glycerol, and then stored at -80°C.

Gifu Anaerobic Medium (GAM, HyServe, cat. no. 5422) was used for downstream analyses, since strain WCA-9-b2^T^ showed improved growth in GAM vs. WCA. GAM agar was supplemented with 2 mL/L titanium(III) chloride (GAM-TC) (Sigma, cat. no. 1107890001) solution [11, 12] as an additional oxygen scavenger to obtain clear colonies on Petri dishes. The anaerobic handling of downstream analyses at UCPH was performed in another anaerobic chamber (Model AALC, Coy Laboratory Products) containing ∼93% (v/v) N_2_, ∼2% H_2_, ∼5% CO_2_ and an atmosphere kept at RT. Agar plates were incubated at 37°C in an anaerobic jar (Cat. No. HP0011A, Thermo Scientific) containing an anaerobic sachet (Cat. No. AN0035A AnaeroGen™, Thermo Scientific) outside the anaerobic chamber.

## 16S rRNA gene phylogeny

Genomic DNA used for 16S rRNA analysis was extracted using the Bead-Beat Micro AX Gravity kit (A&A Biotechnology, cat. no. #106-100 mod.1) following the protocol of the manufacturer. Primers for 16S rRNA gene amplification were 27F 5’-AGA GTT TGA TCC TGG CTC AG-3’ and 1492R 5’-GGT TAC CTT GTT ACG ACT T-3’[13]. Annealing temperature was 54°C and DreamTaq Green was used as DNA polymerase (Thermo Scientific, cat. no. K1081). The nearly complete 16S rRNA gene PCR product was purified and sequenced with Sanger-technology by Macrogen Europe BV by using the two primers aforementioned with the addition of 785F 5’-GGA TTA GAT ACC CTG GTA-3’ and 907R 5’-CCG TCA ATT CMT TTR AGT TT-3’ to ensure sequencing overlap. Low quality nucleotides were removed from the Sanger-sequenced 16S rRNA gene in MEGAX v. 10.0.5 followed by taxonomic identification with EZBioCloud (Update 2019.08.06) [14]. A phylogenetic maximum likelihood tree (bootstrap = 1000) was created to compare against the 30 closest species based on 16S rRNA gene similarity retrieved from EZBioCloud and aligned with ClustalW2 [15]. The 16S rRNA gene from the cyanobacterium *Lyngbya aestuarii* PC 7419 (NR_114680.1) was included as root. Phylogenetic analysis based on a nearly complete 16S rRNA gene sequencing (1481 bp; accession no. MN756014) showed that strain WCA-9-b2^T^ was a member of the family *Lachnospiraceae*, order *Clostridiales. Clostridium scindens* (ATCC 35704) was the closest phylogenetic related bacteria (Fig. 1), but the confidence of branching was low (< 50). WCA-9-b2^T^ was distantly related to genera belonging to family *Lachnospiraceae* (*Dorea, Faecalicatena, Blautia, Coprococcus*), *Ruminococcaceae* (*Ruminococcus*), and *Eubacteriaceae* (*Eubacterium*). The 16S rRNA gene sequence of WCA-9-b2^T^ was ≥ 98% similar to already published sequences of uncultured bacteria isolated from the caecum of both lean (DQ815562[16], JQ084505[17] and EF602954[18]) and obese (EF098864[19]) laboratory mice. All the closest relatives with a validly published taxonomy at EZBioCloud belonged to the order *Clostridiales* as follows: *Dorea longicatena* (AAXB02000032, 94.9% sequence identity), *Ruminococcus gnavus* (AAYG02000025, 94.8%), *Clostridium scindens* (ABFY02000057, 94.3%), *Dorea formicigenerans* (AAXA02000006, 94.2%) *Ruminococcus lactaris* (ABOU02000049, 93.8%), *Clostridium hylemonae* (AB023972, 93.7%), *Merdimonas faecis* (KP966093, 93.7%), *Faecalicatena contorta* (FR749945, 93.5%), *Faecalicatena fissicatena* (LDAQ01000152, 92.9%).

**Fig. 1:**
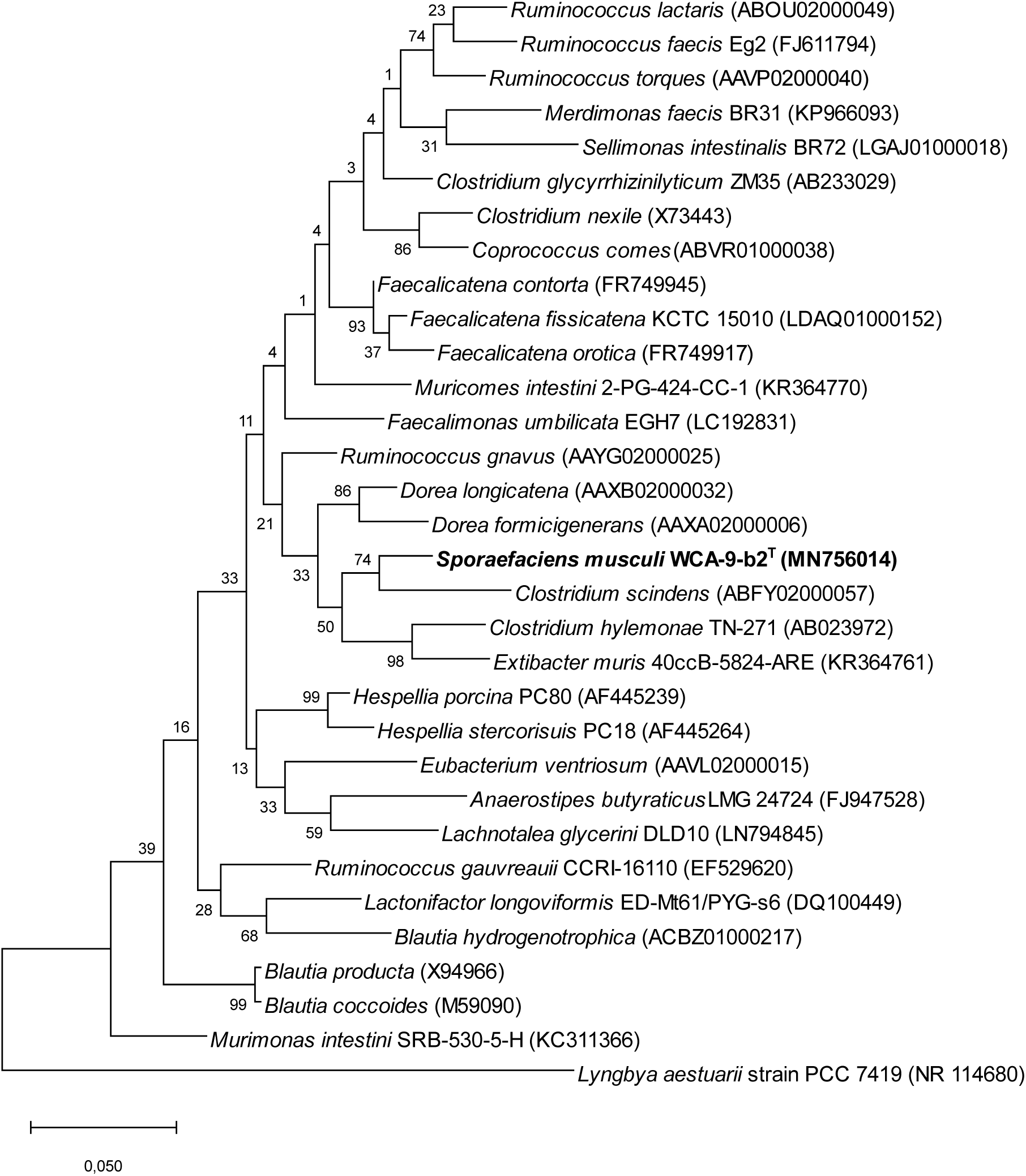
Phylogenetic tree showing the position of Sporaefaciens musculi WCA-9-b2^T^ among the top-30 hits (highest 16S rRNA gene sequence identities) in the EZBioCloud database. 1472 bp (out of 1481 bp) were considered of WCA-9-b2^T^ in the final phylogenetic tree. The GenBank accession numbers of 16S rRNA gene sequences applied to construct the phylogenetic tree are indicated in parentheses. The rooted tree was constructed using the maximum likelihood method and the 16S rRNA gene sequences were aligned with ClustalW2. Cyanobacterium Lyngbya aestuarii was used as an outgroup to root the tree.

## Genomic characterisation of strain WCA-9-b2^T^

For generation of the chromosomal genome, library was constructed and sequenced as previously described [20] with the Illumina NextSeq v2 MID output 2×150 cycles chemistry generating short DNA reads. Additionally, whole-genome sequencing was performed with the MinION platform from Oxford Nanopore Technologies (ONT) to obtain long DNA reads. Genomic DNA was extracted using the MagAttract HMW DNA Kit (QIAGEN, cat. no. 75 3) according to the manufacturer’s instructions for Gram-positive bacteria. DNA was quantified using a Qubit fluorometer (Thermo Fisher Scientific, Waltham, MA, USA) and the genomic library prepared with the Rapid Barcoding Sequencing kit (SKQ-RBK004) from ONT according to manufacturer’s instructions. Sequencing was performed on a FLO-MIN106D R9 flowcell using the MinKNOW software over 72h run (48+24 h). ONT raw FAST5 reads were base called and demultiplexed with Guppy basecaller v.3.3.2 [21] resulting in 2.0 GB of read data. A complete genome was generated by hybrid assembly of the short Illumina and long ONT DNA read sequencing data using the ONT assembly and Illumina polishing pipeline run with Canu v1.8 [22], Racon v1.4.10 [23] and Pillon v1.20 [24]. Raw sequencing Illumina NextSeq (PRJEB35655) and ONT MinION (PRJEB35656) data can be accessed at EMBL-EBI and the ensuing genome assembly at NCBI (PRJNA592877). The genome assembly resulted in three contigs of a total length of 5,763,728 bp: the chromosomal contig spanning 5,426,837 bp; a short contig of 49,487 bp partially overlapping with the chromosomal contig; a putative plasmid (circular) of 287,404 bp. Spore forming bacteria usually exhibits a larger genome size than non-spore formers [3], which would be in accordance with the genome size of strain WCA-9-b2^T^. The chromosomal genome (5,426,837 bp) showed a G+C content at 44.4 % which differed from the phylogenetic related bacteria listed in Table 1. Alignment of the complete genome and the nearly complete 16S rRNA gene suggested that strain WCA-9-b2^T^ contains three rRNA operons. Draft or complete genomes of the nine closest related strains in Table 1 were retrieved from NCBI [25] and included in calculation of the percentage of conserved proteins (POCP) and average nucleotide identity (ANI). POCP values were calculated by following the method described in Qin *et al.* [26] using a genus delineation threshold of 50%. BLASTP (v2.9.0+) [27] was used for protein-protein annotation. POCP analysis of included species (listed in Table 1) provided values < 50%, clearly suggesting that strain WCA-9-b2^T^ represents a novel genus. ANI analysis was additionally performed with OrthoANI [28] and showed ANI values ranging from 69.80 – 74.23% (Table 1), thus supporting that strain WCA-9-b2^T^ represents a novel taxon considering the species-level threshold of 95.0% [29].

**Table 1:** Parameters that differentiate strain WCA-9-b2^T^ from phylogenetically neighbouring taxa. 1 = Sporaefaciens musculi WCA-9-b2^T^, 2 = Dorea longicatena DSM 13814[43], 3 = Ruminococcus gnavus ATCC 29149[44], 4 = Clostridium scindens ATCC 35704[45–47], 5 = Dorea formicigenerans ATCC 27755[43], 6 = Ruminococcus lactaris ATCC 29176[44], 7 = Clostridium hylemonae DSM 15053[46], 8 = Merdimonas faecis strain BR31[48], 9 = Faecalicatena contorta strain 2789S TDY58347876[49], 10 = Faecalicatena fissicatena KCTC 15010[49]. n.r. = not reported, “+” = positive, “-” = negative.

|  | 1 | 2 | 3 | 4 | 5 | 6 | 7 | 8 | 9 | 10 |
| --- | --- | --- | --- | --- | --- | --- | --- | --- | --- | --- |
| Cell shape | Rod | Rod | Cocci | Rod | Rod | Cocci | Rod | Rod | Rod | Rod |
| Spore formation | + | - | + | + | - | - | + | - | - | - |
| α-galactosidase | + | n.r. | + | - | n.r. | n.r. | + | - | + | + |
| β-galactosidase | + | n.r. | n.r. | + | n.r. | n.r. | - | + | + | - |
| α-glucosidase | + | n.r. | n.r. | - | n.r. | n.r. | + | - | + | + |
| α-arabinosidase | + | n.r. | n.r. | n.r. | n.r. | n.r. | n.r. | - | - | - |
| N-acetyl-β-glucosaminidase | + | n.r. | n.r. | - | n.r. | n.r. | - | - | - | - |
| Proline arylamidase | + | n.r. | n.r. | n.r. | n.r. | n.r. | n.r. | - | - | + |
| Sulphide production | - | n.r. | + | + | n.r. | - | n.r. | - | + | + |
| Acetate production | + | + | n.r. | + | + | n.r. | n.r. | + | n.r. | n.r. |
| Succinate production | + | - | n.r. | - | - | n.r. | n.r. | - | n.r. | n.r. |
| ANI (%) | N/A | 72.17 | 70.52 | 74.23 | 72.06 | 69.80 | 71.87 | 72.50 | 69.98 | 69.52 |
| 16S rRNA gene similarity (%) | N/A | 94.91 | 94.77 | 94.28 | 94.15 | 93.81 | 93.70 | 93.69 | 93.53 | 92.91 |
| G+C (mol %) | 44.44 | 41.40 | 42.70 | 46.90 | 40.70 | 42.50 | 48.80 | 47.00 | 46.90 | 45.50 |

Genome annotation was conducted using PROKKA (v1.13.3) [30] and converted into KEGG annotations using PROKKA2KEGG (https://github.com/SilentGene/Bio-py/tree/master/prokka2kegg) for comparison against the KEGG database. CAZy annotation was conducted using diamond blastp [31] considering 40.0% subject and query coverage and a minimum bitscore of 100 against the CAZy database (dbCAN2, v07312019). The genome of WCA-9-b2^T^ contained 5,844 coding sequences, of which 111 were transporters, 16 secretion genes and 662 unique enzymes. Based on KEGG functions, only starch (and thereby glucose) can be utilised as a carbon source which is one of the main components in GAM broth. However, sulphide and L-serine were identified as potential substrates to produce L-cysteine and acetate, which may act as a carbon source (EC:2.3.1.30, 2.5.1.47). The ability to convert acetate to acetyl-CoA may support this energy route (EC:2.3.1.8, 2.7.2.1). Additionally, propionate production was identified from propanoyl-CoA (EC:2.3.1.8, 2.7.2.1). L-glutamate production from ammonia was identified via L-glutamine (EC:6.3.1.2, 1.4.1.) and folate (vitamin B9) biosynthesis from 7,8-dihydrofolate (EC:1.5.1.3). In total, 337 CAZymes were encoded in the genome, including members from 37 glycoside hydrolase families and 11 glycoside transferase families, the largest of which included 92 genes (GT2). A large repertoire of carbohydrate-binding modules was also present, which may suggest this species is specialised to degrade complex carbohydrates. Up to six genes involved in sporulation, including SpmA, SpmB, and GerAA derivatives, as well as 24 genes encoding flagella-related proteins were identified. Antibiotic resistance genes and virulence factors were detected with True ac™ ID [32]. The antibiotic resistance gene *patA* (ARO ID: 3000024) coding for an efflux pump was detected as well as several virulence genes involved in antiphagocytosis (VF0361, VF003, and VF0144), adherence (VF0323), secretion systems (VF0344), and stress protein (VF0074). CRISPRDetect [33] v. 2.1 was applied to predict functional CRISPR-cas systems and classification was based on Makarova *et al.* [34]. A CRISPR-Cas system of strain WCA-9-b2^T^ was suggested to belong to Class II type II-A, based on the composition and ordering of cas-genes (cas9-cas1-cas2-csn2) [34]. BLAST searches demonstrated that none of the 21 spacer sequences matched bacteriophage (phage) DNA available at the NCBI databases or the host genome of WCA-9-b2^T^. It is not yet certain if the detected CRISPR-Cas type II-A system in strain WCA-9-b2^T^ is active. The phage prediction algorithm PHASTER[35] revealed three intact (score > 110) prophages with genome sizes of 22.8 kbp, 30.8 kbp, and 30.3 kbp and phage proteins representing genes encoding terminase, integrase, DNA-packaging, transposase, coat protein, and phage-like proteins with unknown function. Fluctuations in growth curves measured by optical density (OD_600_) indicated potential prophage induction (Fig. S1) and was further supported by a high number of virus-like-particles (VLPs) observed with epifluorescence microscopy (Fig. S2) that was performed as previously described [36].

## Phenotypic characterisation of strain WCA-9-b2^T^

Growth of strain WCA-9-b2^T^ was evaluated by measuring OD_600_ at different pH (5.5, 6.0, 6.5, 7.3, 8.0, 8.5, 9.0, 9.2, 9.5, 10.0) and temperatures (20°C, 30°C, 37°C, 40°C, 42°C, 45°C) in triplicates. Cell morphology and motility of liquid cultures, incubated for 24 hours and 48 hours, were evaluated by phase-contrast microscopy (Olympus BX40 microscope and UPlanFl 100x/1.30 oil immersion objective) and motility was further accessed with incubation in Sulphide Indole Motility [37] (SIM) medium that contained GAM broth powder. ndole production was tested with Kovac’s indole reagent (VWR, cat. no 1.09293.0100). The optimal temperature was 37°C and the optimal pH 7.3; growth was nevertheless observed in the range of 30-40°C and pH 6.5-8.5. Both Gram staining [38] and the potassium hydroxide test (3% (w/v) KOH) [39] showed that the strain is Gram-positive (Fig. S3), which is consistent with most bacteria belonging to the order of *Clostridia*. Strain WCA-9-b2^T^ formed long rods with pointy ends and cell length varies between 1.8 µm – 9.9 µm (average 4.8 µm ± 1.5 µm) and cell width varies between 0.4 µm – 0.8 µm (average 0.6 µm ± 0.1 µm) depending on time of incubation (24-48 hours). Motility of strain WCA-9-b2^T^ was not observed under the conditions tested, despite the presence of numerous flagella genes in the genome. The SIM test was also negative for sulphide and indole production. Two weeks-old liquid cultures were subjected to endospore staining following the Schaeffer-Fulton protocol [40] and observed in bright-field with 1000x magnification. Although it was a rare event, Schaeffer-Fulton staining revealed endospore and spore formation (Fig. S4). For scanning electron microscopy (SEM), the strain was grown in GAM medium until the exponential phase was reached after 30 hours with bacterial density nearing McFarland standard 4. Bacteria were placed at a 0.2 µm polycarbonate filter as follows. A small amount of sterile water was added to a vacuum slot. On the meniscus of the water a filter paper (Whatman, cat. no. 1822 025) and a polycarbonate filter (Osmonics, cat. no. 11013) were placed. By applying vacuum excess water was removed and the two filters were brought in close contact. Vacuum was released and 10 µl of bacterial culture was placed in the middle of the polycarbonate filter. The suction from the underlying Whatmann filter gently removed the water. Immediately when the drop of water had disappeared the polycarbonate filter was transferred to Karnovsky’s fixative. n the following fixation and dehydration procedure the filters were placed floating on the surface of the liquids with bacteria on the upper side. Further fixation was obtained with osmium tetroxide, followed by a graded series of ethanol, and final drying with hexamethyldisilazane. The cells were subsequently mounted to an aluminium stub, coated with gold-palladium and observed in a Quanta 200 SEM at 15kV. No flagella-like elements were observed by SEM (Fig. 2), supporting the findings above. In contrary, SEM imaging identified spore-like formation (Fig. 2a). The SEM imaging was in line with the Schaeffer-Fulton staining, annotated genes involved in sporulation and the initial spore selecting procedure of the caecal content, which overall confirmed the spore forming ability of strain WCA-9-b2^T^.

**Fig. 2:**
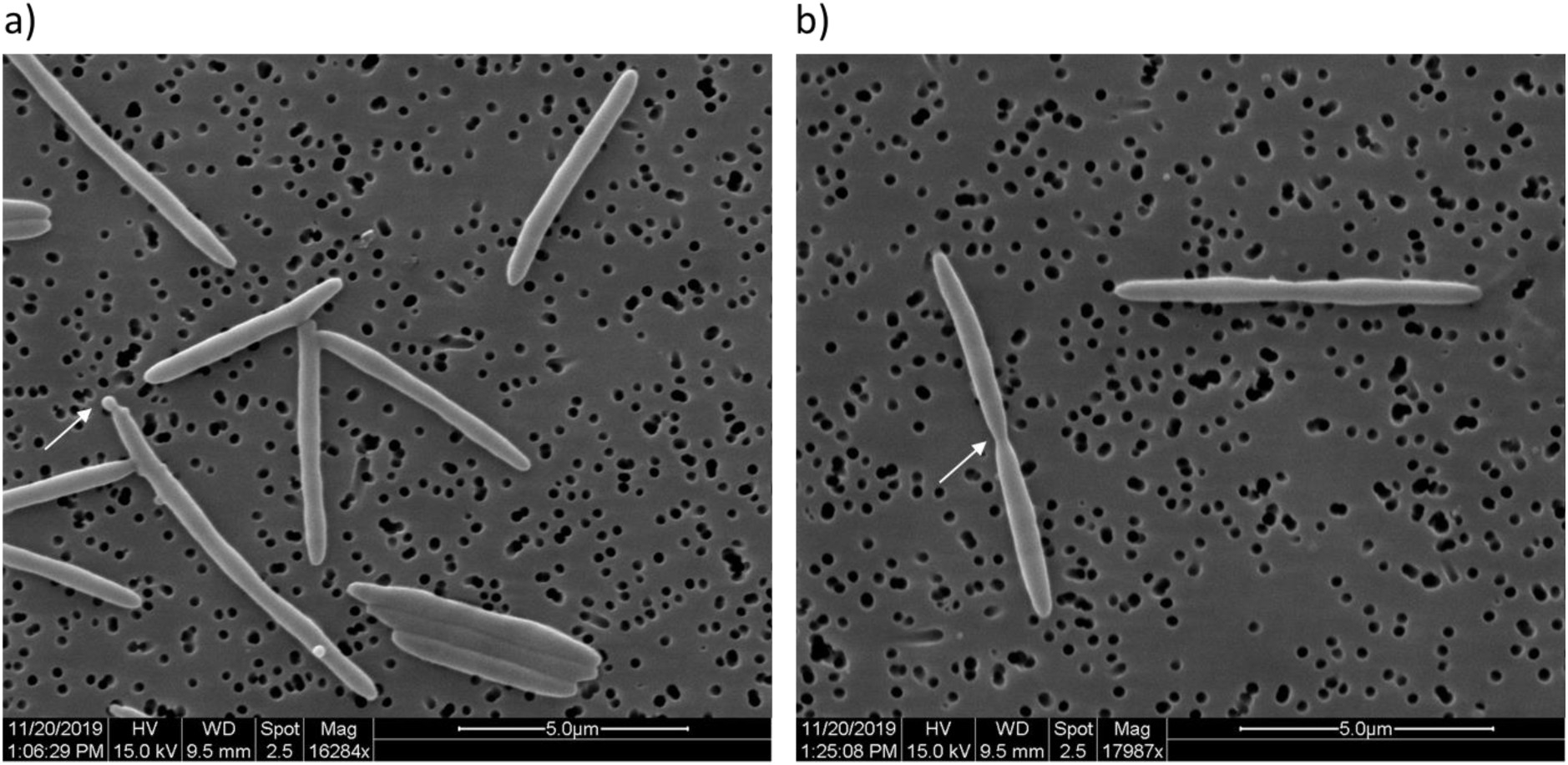
Scanning electron microscopy (SEM) images of strain WCA-9-b2^T^. Flagella were not observed. a) Spore formation was observed at a low frequency (white arrow). b) one cell is at the final stage of division.

Enzymatic activities were determined in triplicates using API® Rapid ID 32A strips following the manufacturer’s instructions (iomérieux, cat. no. 32300, 70700, 70 40, 70542, 70442, 70562, 70100, and 70900). The enzymatic assay demonstrated that the strain WCA-9-b2^T^ was positive for α- and β-galactosidase, α-glucosidase, α-arabinosedase, N-acetyl-β-glucosaminidase, and proline arylamidase. The enzymatic profile of WCA-9-b2^T^ had no match with the identification table of Rapid ID32 A v3.2.

The concentration of SCFAs (acetate, propionate, butyrate, valerate), branched SCFAs (isobutyrate, isovalerate), and intermediate metabolites (lactate, succinate, formate) were determined using high-performance liquid chromatography refractive index (HPLC-RI). Bacteria were grown in modified YCFA broth (DSMZ medium 1611) supplemented with 0.02 % DTT in Hungate tubes for 48h at 37 °C under anaerobic conditions (89.3% N_2_, 6% CO_2_, 4.7% H_2_) and with shaking (200 rpm). The strain was grown in triplicates and the negative control consisted of medium without bacteria. After incubation, samples were centrifuged (10,000 g, 10 min, RT) and supernatants were collected and stored at -20 °C until analysis. Before HPLC-RI analysis, samples were filtered into 2 ml short thread vials with screw caps (VWR International GmbH, Germany) using non-sterile 0.2 µm regenerated cellulose membrane filters (Phenomenex, Germany). Vials were then placed in the refrigerated autosampler of the HPLC system, a Hitachi Chromaster 5450 (VWR International GmbH, Germany) fitted with a Refractive Index detector and a Shodex SUGAR SH1011 column (300 x 8.0 mm) (Showa Denko Europe, Germany). A Shodex SUGAR SH-G (6.0 x 50 mm) was used as guard column. The injection volume was 40 µl. The oven temperature was 40 °C. The eluent was 10 mM H_2_SO_4_ with a constant flow of 0.6 ml/min. Concentrations were determined using external standards via comparison of the retention time (all compounds were purchased from Sigma-Aldrich). Peaks were integrated using the Chromaster System Manager software (Version 2.0, Hitachi High-Tech Science Corporation). For each of the tested strains, only SCFA concentrations >0.8 mM (limit of detection for succinate, lactate, and acetate) or >0.5 mM (LOD for glucose and all other SCFAs) in at least one of the triplicates were considered for calculation.

HPLC-RI analysis showed that WCA-9-b2^T^ metabolised glucose (−4.4 mM), which agrees with the α-glucosidase activity measured by enzymatic tests. Succinate (5.7 mM) and acetate (12.9 mM) were produced under the experimental conditions tested. The production of succinate and acetate suggest that strain WCA-9-b2^T^ is involved in cross-feeding other bacteria in the gut which are able to convert succinate and acetate into the SCFAs butyrate or propionate [41].

The IMNGS platform[42] was used to screen for the relative abundance of WCA-9-b2^T^ (> 97% similarity) in 4721 samples from various studies investigating the bacterial GM component of mice. The relative abundance of bacteria sharing more than > 97% rRNA gene identity with WCA-9-b2^T^ was below 0.08% (representing > 500 sequencing read counts), and was found in both lean (SRR1698289, ERR1173727), obese (SRR2073437, SRR1959789) and antibiotic treated (SRR1960027) mice. Although strain WCA-9-b2^T^ was isolated from an obese mouse, there is currently no evidence that its occurrence is associated with the host phenotype.

Based on all the aforementioned genotypic and phenotypic characteristics, we suggest that strain WCA-9-b2^T^ should be designated the type strain of a novel bacterial genus and species within the family *Lachnospiraceae*, order *Clostridiales*, for which the name *Sporaefaciens musculi* is proposed. Parameters that help distinguishing *S. musculi* WCA-9-b2^T^ from phylogenetically most closely related bacteria are listed in Table 1:

### Description of *Sporaefaciens gen. nov*

*Sporaefaciens* (Spo.rae.fa’ci.ens. NL. N. *sporae*, spore; L. masc. n. *faciens*, making; N.L. masc. n. *Sporaefaciens*, spore-maker)

Bacteria of the genus *Sporaefaciens* are strictly anaerobic, spore-forming, Gram-positive rods. Motility has not yet been observed. Based on 16S rRNA gene and genome analysis, the genus *Sporaefaciens* belongs to family *Lachnospiraceae* (phylum *Firmicutes*, order *Clostridiales*) is distantly related (≤ 95% identity between 1 S rRNA gene sequences; POCP values <50%) to other genera members belonging to the such as *Clostridium, Dorea, Ruminococcus, Merdimonas, Extibacter*, and *Faecalimonas.* The type species is *Sporaefaciens musculi*.

### Description of *Sporaefaciens musculi gen. nov., sp. nov*

*Sporaefaciens musculi* (mus’cu.li. L. gen. n. *musculi*, of a common mouse)

The species possesses all the features of the genus. Cultures in the stationary phase appear with turbidity resembling McFarland standard 4. Cell length varies between 1.8 µm – 9.9 µm (average 4.8 µm ± 1.5 µm) and cell width varies between 0.4 µm – 0.8 µm (average 0.6 µm ± 0.1 µm). Cells appear as single cells or in pairs. After 72 hours on GAM-TC agar, colonies are transparent with irregular shape and a diameter of 1-2 mm. Optimal growth occurs at 37°C and pH 7.3. In modified YCFA, the species was able to use glucose and produced succinate and acetate. Using the API® Rapid ID 32A test, strain WCA-9-b2^T^ is positive for α- and β-galactosidase, α-glucosidase, α-arabinosedase, N-acetyl-β-glucosaminidase, and proline arylamidase. It is negative for urease, arginine dihydrolase, β-galactosidase 6 phosphate, β-glucosidase, β-glucuronidase, glutamic acid decarboxylase, α-fucosidase, alkaline phosphatase, arginine arylamidase, leucyl glycine arylamidase, phenylalanine arylamidase, leucine arylamidase, pyroglutamic acid arylamidase, tyrosine arylamidase, alanine arylamidase, glycine arylamidase, hisitidine arylamidase, glutamyl glutamic acid arylamidase, serine arylamidase, indole production, as well as mannose and raffinose fermentation and reduction of nitrates. Its G+C content of the genomic DNA is 44.4%. The type strain is WCA-9-b2^T^ (=DSM 106039^T^ = CCUG (pending ID^T^). It was isolated from the caecal content of an 18-week-old male C57BL/6NTac mouse fed a high-fat diet for 13 weeks.

## Abbreviations

WCA: Wilkens-Chalgren-Anaerobe;
GAM: Gifu Anaerobic Medium;
RT: Room Temperature;
ONT: Oxford Nanopore Technologies;
ANI: Average Nucleotide Identity;
SEM: Scanning Electron Microscopy;
SIM: Sulphide Indole Motility;
HPLC-RI: High Performance Liquid Chromatography – Refractive Index;
SCFA: Short Chain Fatty Acid;
CRISPR: Clustered Regularly Interspaced Short Palindromic Repeats;
VLPs: virus-like-particles;
TUM: Technical University of Munich;
UPCH: University of Copenhagen;
POCP: Percentage of conserved proteins.

## Acknowledgement

We thank Andreas Czempiel (ZIEL Core Facility Microbiome, Technical University of Munich) for training and technical assistance during bacterial isolation.

## Ethical statement

Experimental work with mice was carried out in accordance with the Directive 2010/63/EU and the Danish Animal Experimentation Act (licence ID: 2012-15-2934-00256).

