## Supplemental materials and figures for "*Sporaefaciens musculi* gen. nov., sp. nov., a novel bacterium isolated from the caecum of an obese mouse"

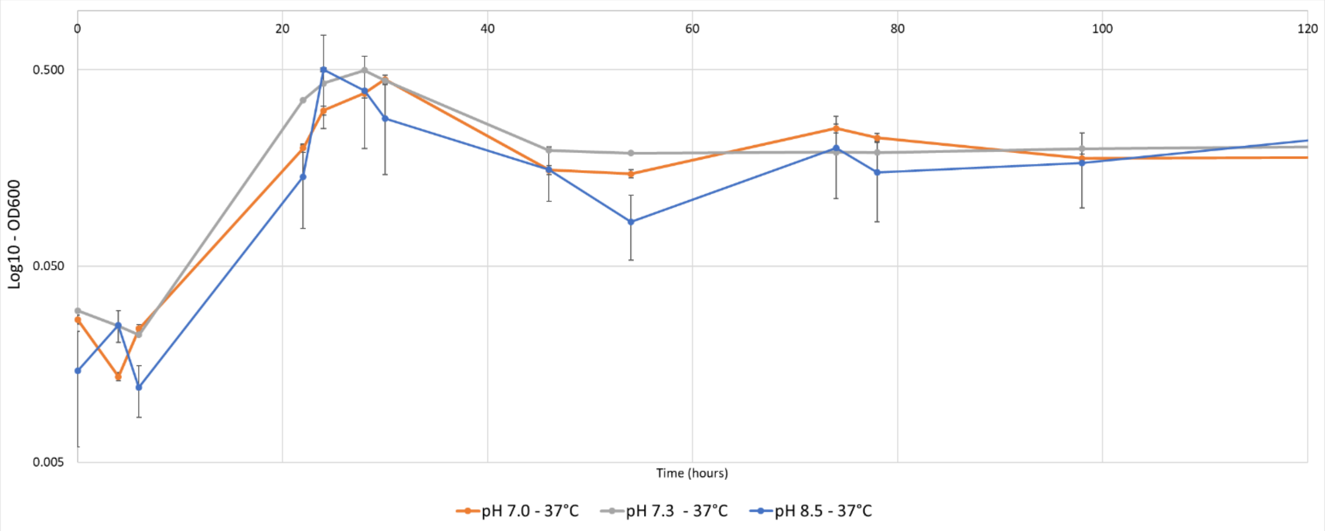


*Fig. S1: Growth curves of strain WCA-9-b2$^{T}$ based on $OD_{600}$ measurements. The cultures were grown in GAM broth at pH 7.0 (orange), pH 7.3 (grey), and pH 8.5 (blue) at 37°C. The drops in optical density starting at ~22 hours and last until ~50 hours may indicate an induction of prophages.*

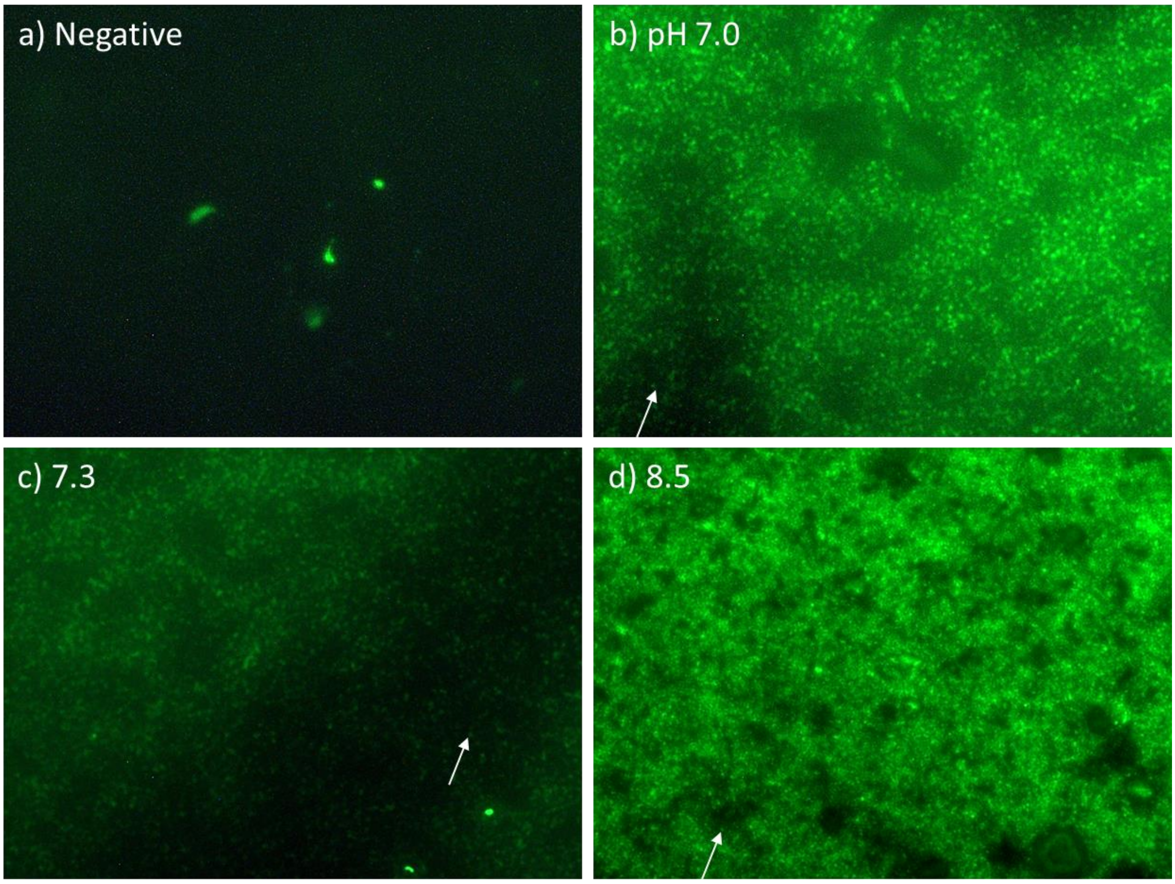


*Fig. S2: Epifluorescence microscopy images of filter-sterilized and 50x concentrated supernatant of strain WCA-9-b2$^T$ grown in GAM broth at pH 7.0, pH 7.3, and pH 8.5 at 37°C (same cultures as in Fig. S1). These images indicated the presence of virus-like particles (VLPs) that might reflect induced prophages of which is marked with white arrows. a) Control (GAM broth), b) strain WCA-9-b2$^T$ at pH 7.0, c) strain WCA-9-b2$^T$ at pH 7.3, d) strain WCA-9-b2$^T$ at pH 8.5.*

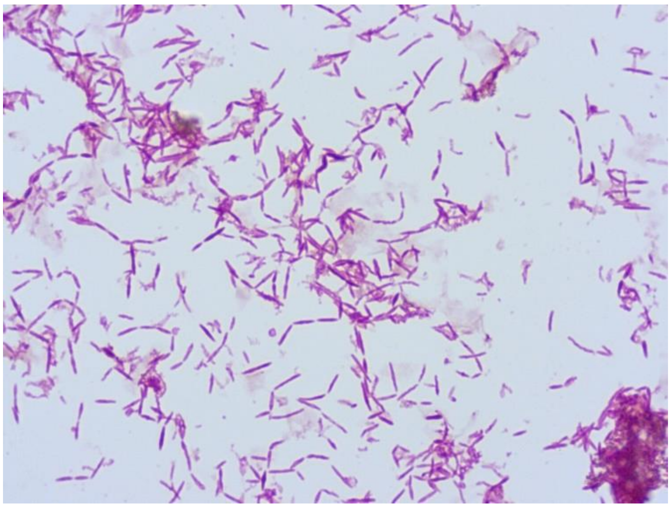

*Fig. S3: Gram-staining of strain WCA-9-b2$^T$ was Gram-positive.*

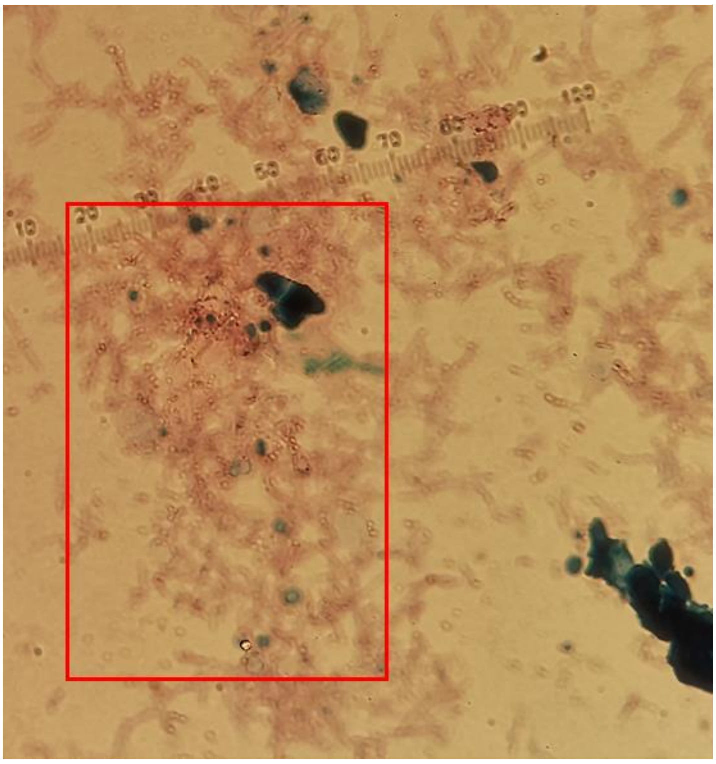

*Fig. S4: Bacterial spores of strain WCA-9-b2$^{T}$ stained with the Schaeffer-Fulton protocol. The red square indicates an area with stained spores.*
